# The key lethal effect existed in the antibacterial behavior of short, medium, and long chain fatty acid monoglycerides on *Escherichia coli*

**DOI:** 10.1101/339309

**Authors:** Song Zhang, Jian Xiong, Wenyong Lou, Zhengxiang Ning, Denghui Zhang, Jiguo Yang

## Abstract

Monoglyceride is an amphiphilic molecule with promising antimicrobial activity for bacteria; however, the key lethal effect in its antibacterial behavior was still unknown. In the study, monobutyrate (MB), monolaurate (ML), monomyristate (MM) were selected to represent the short, medium, and long chain monoglycerides to compare their inhibitory effect against *Escherichia coli*, and a new dose-dependent inhibitory mechanism was proposed by the key lethal effect. The minimal inhibitory concentration and antibacterial curve showed a huge diversity existed in biology activity of MB, ML and MM. The results in scanning electron microscopy and flow cytometry assay indicated that the interference level of MB on cell membrane was obviously weaker than that of ML and MM, while the latter two had similar performance in increasing cell permeability at low doses. The results presented in UV-Vis spectroscopy, cell cycle and biomacromolecules synthesis inhibition assay showed that the cell cycle of *Escherichia coli* was obviously affected by three monoglycerides at doses near MIC, which was therefore regarded as the key lethal effect. The reason for the better biological activity of MM than ML was the stronger interference ability on bacterial cell cycle. In addition, an expanded antibacterial mode was raised that cell permeability increase at low doses was antimicrobial basis, cell cycle arrest at medium doses played the key lethal effect, and cell lysis at high doses was the result of combined action.

## 1. Introduction

*Escherichia coli O157:H7 (E. coli O157:H7)*, a common foodborne pathogenic bacterium, causes a series of infectious diseases in human body such as hemorrhagic diarrhea and enteritis, hemolytic uremic syndrome and thrombotic thrombocytopenic purpura (1–3). This bacteria has a strong pathogenicity and can cause obvious symptom at a very low infection dose which is lower than 50—100 cells (4). It was firstly discovered and isolated from the feces of food poisoning patients consuming contaminated hamburger in America, and subsequently appeared some large-scale outbreak cases in Japan, Canada, Australia, and some Nordic countries (4–8). The most serious disease outbreak occurred in Japan, and the infected population was up to tens of thousands of people, causing 11 of death (9). According to the report from The Centers for Disease Control, *E. coli O157:H7* infection caused 20000 disease cases and 200—500 death each year with estimated annually medical expenses of 405 million dollars in the United States (10, 11). This toxin-producing pathogenic *E. coli* has become the second common intestinal pathogen following Salmonella, and *E. coli O157:H7* infection accounted for 91.4% in pathogenic *E. coli* diseases (12).

Microbial contamination has become a major source of safety risks in food industry, and various physical treatments and chemical additives have been used to eliminate potential pathogens (13, 14). Owing to the concern about safeness of chemicals, consumers prefer natural preservatives or bactericides isolated from some animal and plant materials instead of synthetic chemicals (15). Fatty acid monoglycerides are a class of promising antibacterial agents, naturally occurred in American pusa, animal milk and some other foods (16, 17). The esters have broad antibacterial activity toward gram-negative, gram-positive bacteria and fungi including yeasts and moulds, and inhibition effects are little affected by pH (18). In addition, monoglycerides can be resolved in the gastrointestinal tract and do not show any toxicity to human body (19). In the past decade, the studies on antibacterial activity of monoglycerides have been limited to some medium carbon chain glycerides ranged from 8 to 12 carbons (20–23). Recently, some literatures reported that the antimicrobial activity of long carbon chain fatty acid monoglycerides, including monomyrisate and monopalmitate (24, 25). Clelia Altieri revealed that the inhibition indexes of monomyrisate and monopalmitate were obviously higher than that of monolaurin on *E. coli O157:H7* at 20 ppm after 10 h of incubation, and the inhibitory effectiveness was in dose-dependent for monolaurin, but the two else were exactly the opposite (26). However, the systematic studies on the antibacterial effect of short, medium and long carbon chain fatty acids monoglycerides are still lacking, more comparative studies are therefore necessary if selecting suitable monoglycerides for the inhibition of different microorganisms.

The action mechanisms in most monoglycerides antimicrobial studies focused on membrane damage, which could increase cell permeability and even lead to the leakage of cellular contents (17, 21, 27). Hyldgaard, M. visualized membrane disruption caused by monocaprylate using atomic force microscopy, indicating that cell membrane was a important action site in antibacterial test (28). Although it is known that monoglyceride play the inhibitory effect by membrane damage, the complete antimicrobial mechanism is still not fully understood. Interestingly, the cell wall of the bacteria with incomplete cell membrane gradually split in the late stage of antibacterial test, suggesting that the cell wall lysis was obviously later than the membrane disruption, which was more like an antibacterial result than a action mode (29). Recently, scientists also found some cationic antibacterial peptides not only formed holes on membranes, but also bound to DNA and interfered normal cell metabolism, indicating that intracellular action was a non-negligible process after antimicrobial agents penetrating through cell membrane (30–32). Therefore, the potential intracellular action goals, especially for DNA, should be incorporated into research scope in antibacterial test in the future.

In present study, the inhibitory effect of short chain (2 — 6 carbons), medium chain (8 — 12 carbons), and long chain (14—18 carbons) fatty acid monoglycerides on *E. coli O157:H7* were compared by minimal inhibitory concentration (MIC) and antibacterial curves. The effects of monoglycerides with different chain lengths on cell surface and membrane permeability were evaluated by scanning electron microscopy (SEM) and flow cytometry. In addition, the interaction between monoglyceride and genomic DNA were studied by UV-visible spectrum and cell cycle. Moreover, intracellular DNA, RNA and protein detection was employed to explain the possible relation between DNA double helix disruption and cell division inhibition. This study was aimed at seeking the key lethal effect in monoglyceride antibacterial test and to explain the reason for the difference in the antibacterial effect of short, medium, and long chain monoglycerides.

## 2. Materials and methods

### 2.1 Materials

Monoacetate (MA), monobutyrate (MB), monocaprylate (MC), monolaurate (ML), monomyristate (MM) and Monopalmitate (MP) were obtained from Molbase Chemical Co. (Shanghai, China) with purity≥99.0%. A series of stock solutions of the six monoglycerides were made in ethanol to obtain the concentrations of 0.01, 0.02, 0.04, 0.08, 0.16, 0.32, 0.64, 1.25, 2.5, 5, and 10 mg/mL, respectively. Propidium iodide (PI, purity≥94.0% in HPLC) was purchased from Sigma-Aldrich Co. (Santa Clara, USA). A PI stock solution was prepared to achieve the concentration of 10 mg/mL in phosphate buffer solution (PBS, 0.1 M, pH 7.5, Sigma-Aldrich Co., Santa Clara, USA) and stored at 2-8°C. Hoechst 33342 (HO, purity≥98%) and fluorescein isothiocyanate (FITC, purity≥90%) were purchased from Yuanye Biological Technology Co. (Shanghai, China), while Pyronin Y (PY, high purity biological stain) was obtained from Acros Co. (Belgium). Their stock solutions were prepared in PBS to harvest 500 μg/mL of HO, 2 mg/mL of PY, and 100 μg/mL of FITC. An ezup column bacteria genomic DNA purification kit was bought from Sangon Biotech Co. (Shanghai, China). Tris-HCl buffer (0.05 M, pH 7.4) was obtained from Yuanye Biological Technology Co. (Shanghai, China). The water used in the study was purified by Milli-Q device (Merck Millipore Co., Massachusetts, USA). Glutaraldehyde solution (25% in H_2_O) and osmium tetroxide (purity≥98%) were bought from Sigma-Aldrich Co. (Santa Clara, USA). 2.5% (v/v) glutaraldehyde work solution and 2% (w/v) osmium tetroxide work solution were prepared in PBS for later use.

### 2.2 Experimental strain

A gram-negative bacterium, *E. coli O157:H7* was purchased from Guangdong Culture Collection Center (Guangzhou, China). The strain powder was activitied by dissolving with 1ml sterile PBS and then transferred to tryptone soy agar (TSA, Shuoheng Biotechnology Co., Guangzhou, China) solid plate to culture for 24 h or longer at 37°C. Subsequently, a single colony on solid medium was hooked onto the TSA slope medium with multiple scribing operations and cultured for 24—48 h before storing at 2—8°C.

### 2.3 Antibacterial activity assays

The antibacterial effects of short, medium and long carbon chain fatty acid monoglycerides on *E. coli* were characterized by detecting MIC and inhibition curves (33, 34). The refrigerated strain was cultured in 100 mL tryptic soy broth (TSB, Shuoheng Biotechnology Co., Guangzhou, China) medium with 120 rpm shaking in 37°C until mid-logarithmic growth phase and centrifugated at 3000 rpm for 5 min. The cell pellets were firstly washed with PBS and then resuspended into diluent with cell concentration of approximate 10^5^ CFU/mL in TSB, which was corresponding to optical density of 0.3 at 600 nm (OD_600_≈0.3). 20 μL of different concentrations of MA, MB, MC, ML, MM and MP stock solutions were blended with 180 μL of *E. coli* diluent to achieve final concentrations of 1, 2, 4, 8, 16, 32, 64, 125, 250, 500, and 1000 *μg/mL* in 96 well plates. The controls were adding the same volume of ethanol, and each concentration was repeatly performed three times. The plates were cultured at 37°C for 24 h with 120 rpm shaking. The OD_600_ values were recorded at 0 h and 24 h by a microplate reader (SpectraMax i3x, Molecular Devices Co., San Jose, USA). The MIC was defined as the lowest concentration in which the OD_600_ increase was smaller than 0.05 during 24 h.

The inhibition curves reflected the vitality of bacterial cells in monoglyceride treatment and were measured by plate counting method. According to the results of MIC assay, MB, ML and MM were selected for representing short, medium and long chain fatty acid monoglycerides to complete the research. 50 μL of MB, ML, and MM solutions were mixed with 950 μL of *E. coli* diluent in sterile 1 mL centrifuge tubes to reach final concentrations of 8, 16, 32, 64, 125, 250, 500, and 1000 μg/mL, respectively. The controls were adding 50 μL of ethanol and all samples were conducted in triplicate. The cells were cultivated for 1 h at 37°C with 120 rpm shaking. Then 10-fold serially dilutions were performed for all *E. coli* samples and cultivating for 24 h at 37°C prior to colony counting.

### 2.4 Membrane integrity test

#### 2.4.1 SEM assay

According to the reports of previous literatures (35, 36), the influence of MB, ML and MM on the *E. coli* surface was assessed by SEM. 50 μL of three monoglycerides solutions were mixed with 950 μL of cell diluent prepared as the above method in sterile centrifuge tubes to obtain final concentrations of 1/2 MIC, 1 MIC, and 2 MIC, respectively. The controls were adding the same volume of ethanol. The cell samples were cultured for 1 h at 37°C with orbital shaking at 120 rpm and collected by centrifugation at 3000 rpm for 5 min. Then cell pellets were washed twice with sterile PBS and fixed with 0.5 mL of 2.5% (v/v) glutaraldehyde work solution for 12 h at 4°C. On the second day, the *E. coli* cells were post fixed in 0.1 mL of 2% (w/v) osmium tetroxide work solution at room temperature for 2 h. Subsequently, the immobilized cells were washed with sterile PBS and orderly dehydrated with 30%, 50%, 60%, 70%, 80%, 90%, and 100% ethanol solution for 10 min. Then a drop of cell solution in ethanol was added onto the coverslip to stand until the cells had settled down. After freeze-drying in vacuum for 24h or longer, a cold field scanning electron microscopy (UHR FE-SEM SU8220, Hitachi Ltd., Tokyo, Japan) was allowed to record the change of membrane morphology.

#### 2.4.2 Cell permeability measurement

The changes of *E. coli* membrane permeabilities after adding MB, ML and MM were investigated by the method of Hyldgaard, M. et al. with some modification (37). The *E. coli* cells were harvested in mid-log growth phase by centrifugation and wash, and resuspended to cell density of 10^5^ CFU/mL with sterile TSB. 950 μL of aliquot cell diluents were combined with 50 μL of three monoglycerides solutions to obtain final concentrations of 0, 2, 4, 8, 16, 32, 64, 125, 250, and 500 μg/mL. Each concentration was performed three times in parallel. All groups were cultured at 37°C for 1 h except of 85°C heating for 1 h in positive controls, which provided maximum detection boundary (MDB). Subsequently, the cells were washed with sterile PBS and resuspended in 1 mL PBS before staining with 50 μL of 100 μg/mL PI work solution for 20 min in dark condition. Finally, the cell samples were immediately allowed to fluorescence analysis by a flow cytometer (CytoFLEX, Beckman Coulter Co., California, USA). The excitation and emission wavelength of PI were located at 488 nm and 610 nm, respectively (38). The sample flow rate was controlled as 100—500 cells/s, and at least 10000 cells were collected for the following data analysis.

### 2.5 Interaction of monoglycerides with different chain lengths and genomic DNA

#### 2.5.1 Impact of different monoglycerides on the structure of genomic DNA

To examine the impact for DNA brought by monoglycerides, UV-visible spectroscopy was carried out similar to the method described by Hegde, A. H. et al (39). The genomic DNA was purified from viable *E. coli* cells in mid-log growth phase by an ezup column bacteria genomic DNA purification kit. The ratio of the UV absorbance of genomic DNA at 260 nm and 280 nm was 1.85, suggesting that the purified DNA was free from proteins (40). The concentration of *E. coli* DNA was diluted to 3.6 mM in sterile Tris-HCl buffer, calculating by UV absorbance at 260 nm in 1 cm quartz cell divided by 6600 M^−1^cm^−1^ of a molar absorption coefficient (32). Subsequently, 25 μL of MB, ML and MM solution were mixed with 475 μL of *E. coli* diluents with cell density of 10^5^ CFU/mL to obtain final monoglycerides concentrations of 2, 4, 8, 16, 32, 64, 125, 250, 500, and 1000 μg/mL in sterile 1.5 mL centrifuge tubes. The control groups were adding the same volume of ethanol and all samples were allowed to equilibrate for 5 min before UV spectral scanning. To eliminate the adverse effect from background, the baseline was firstly corrected for Tris-HCl buffer signal. The UV spectral was recorded at a wavelength range from 220 nm to 320 nm and obtained the average of three determinations in parallel.

#### 2.5.2 Effect of different monoglycerides on the cell division of *E. coli*

Cell cycle is an important indicator of cell division function and is measured by flow cytometry combined with PI staining according to the report from Steen, H. B. et al (41). 50 μL of MB, ML and MM solutions were combined with 950 μL of *E. coli* suspensions (10^5^ CFU/ml) to achieve final monoglyceride concentrations of 1/4 MIC, 1/2 MIC, 1 MIC, 2 MIC, and 4 MIC, respectively. The controls were added 50 μL of ethanol and all groups were repeatly performed three times. The treated cells were cultured at 37°C for 1 h with 120 rpm shaking before centrifuge collection and PBS wash. The cell pellets were fixed with 70% (v/v) ice ethanol (pre-cooling at −20°C overnight) for 12 h or longer at 4°C. Subsequently, the cells were centrifuged and washed to remove the stationary liquid. Finally, 1 mL of 50 μg/mL PI work solution (containing 1 mg/mL Rnase) was added into the immobilized cells to stain for 20 min at 4°C in the dark condition before detecting by the Beckman flow cytometer. The excitation and emission wavelength of PI-DNA complex were located at 488 nm and 610 nm, respectively. The flow rate of cell solution was turned to 100—200 cells/s, and at least 30000 cells were captured for the subsequent scatter and histogram analysis.

#### 2.5.3 Influence of different monoglycerides on the synthesis of intracellular biomacromolecules

The synthesis inhibition of MB, ML and MM on DNA, RNA and protein in *E. coli* was studied by flow cytometry combined with three fluorescence staining (42). Three dyes, HO, PY and FITC, were used to stain intracellular DNA, RNA and protein, respectively. Their relative contents were characterized by measuring light intensity of blue, red and green fluorescence which almost had no overlap in the emission spectrum region. The viable *E. coli* cells were harvested from overnight cultures by centrifugation and resuspended to cell density of 10^5^ CFU/mL in broth. 950 μL aliquots of cell suspensions were supplemented with 50 μL of MB, ML and MM solution to reach final concentration of 1 MIC. The controls contained 50 μL of ethanol instead of monoglycerides solutions. The solutions were incubated for 0, 10, 20, 30, 40, 50, and 60 min at 37°C, respectively. After different time periods, the *E. coli* cells were collected by centrifugation. Subsequently, 1 mL of 70% (v/v) ice ethanol (pre-cooled at −20°C for 12h) was added to fix overnight at 4°C. The stationary liquid was removed by centrifugation prior to PBS wash. Finally, 1 mL of mixed dyes solution containing 0.5 μg/mL HO, 2.0 μg/mL PY, and 0.1 μg/mL FITC in PBS was added to stain for 20 min at 4°C in the dark condition before allowing to the analysis of flow cytometry.

The semi-automatic device was equipped with three laser excitation flow system. Specifically, the excitation laser wavelengths of DNA, RNA and protein fluorescence were located at 355 nm, 530 nm, and 457 nm respectively. Correspondingly, the emission fluorescence of HO-DNA (blue), PY-RNA (red), and FITC-protein (green) were measured at 450 nm, 580 nm, and 520 nm, respectively. The fluorescence signal was repeatly captured three times for each cell sample before drawing frequency distribution histograms and calculating phase proportions.

### 2.6 Statistical analysis

The averages of colony count of three repetitions were converted into cell density (log CFU/mL). All data in figure were showed as average ± standard deviation (SD) of three replicate measurements. The variance analysis of results was performed by One-Way ANOVA in OriginPro 8.5. A tukey test was used to confirm the significant variance in statistics between controls and samples when p<0.05.

## 3. Results

### 3.1 Antibacterial effects of monoglycerides with different chain lengths

The antimicrobial activity of short, medium, and long chain monoglycerides was qualitatively compared by MIC and quantitatively characterized by antibacterial curve. The MIC of MB, ML, and MM against *E. coli* were 500 μg/mL, 64 μg/mL, and 32 μg/mL, suggesting that ML and MM are more active than MB in terms of inhibitory concentration. Subsequently, the population of *E. coli* cells after three monoglycerides treatment was shown in Figure 1. The cell counts all decreased in a concentration-dependent manner with the larger decline in ML and MM groups. Besides, the cell population declined more than 2 log units when adding MB, ML, and MM at respective MIC, which exceeded the action effect of three monoglycerides at corresponding half maximal inhibitory concentrations. Furthermore, 500 μg/mL of ML and 250 μg/mL of MM played totally bactericidal activity against *E. coli*, which could not achieve for MB even if its concentration increased to 1000 μg/mL, suggesting that evident variance was existed in the sensitivity of *E. coli* to monoglycerides with different chain lengths.

**Figure 1.**
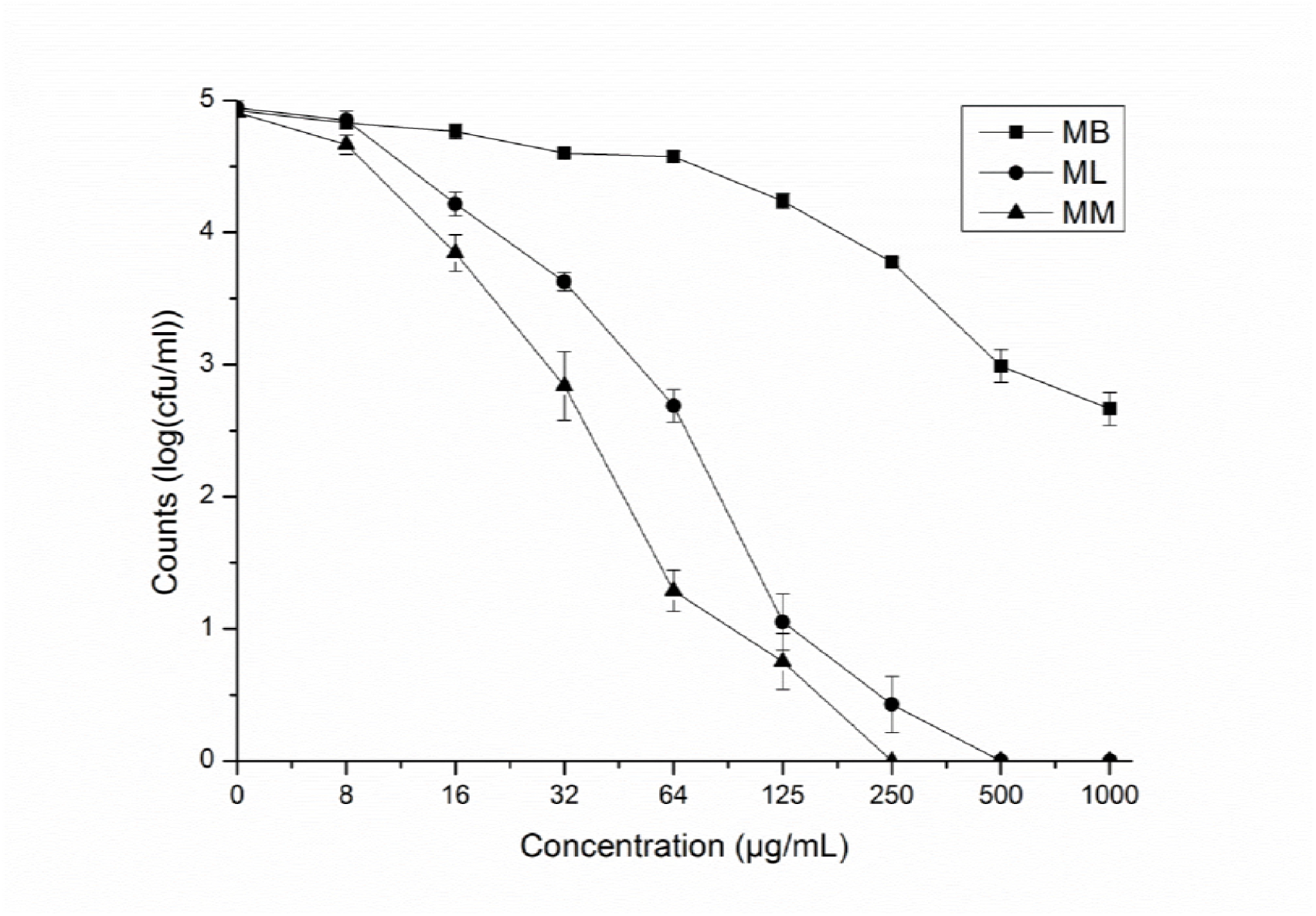
Counts of viable *E. coli* cells after treatment with MB, ML, and MM at 8, 16, 32, 64, 125, 250, 500, and 1000 μg/mL for 1 h. Error bar was represented for standard deviations (SD) of three repeatly determinations (n=3).

### 3.2 Membrane integrity assay

#### 3.2.1 SEM observation

As shown in Figure 2A, the untreated *E. coli* cells had a flat, smooth surface. However, after adding MB, ML, and MM, the cell membrane became uneven, rough, and even concave. Specifically, the *E. coli* cells did not change into rough until MB concentration increased to 2 MIC (Figure 2B-D). After ML treatment, the cell membrane lose smooth and flat surface at 1/2 MIC, and increasing concentration further to 1 MIC and 2 MIC, the cell surface showed further distortion with deeper depression and larger deformation. As for MM treatment at 1/2 MIC, 1 MIC, and 2 MIC, the cells surface sequentially appeared flat, rough, and obvious depression.

**Figure 2.**
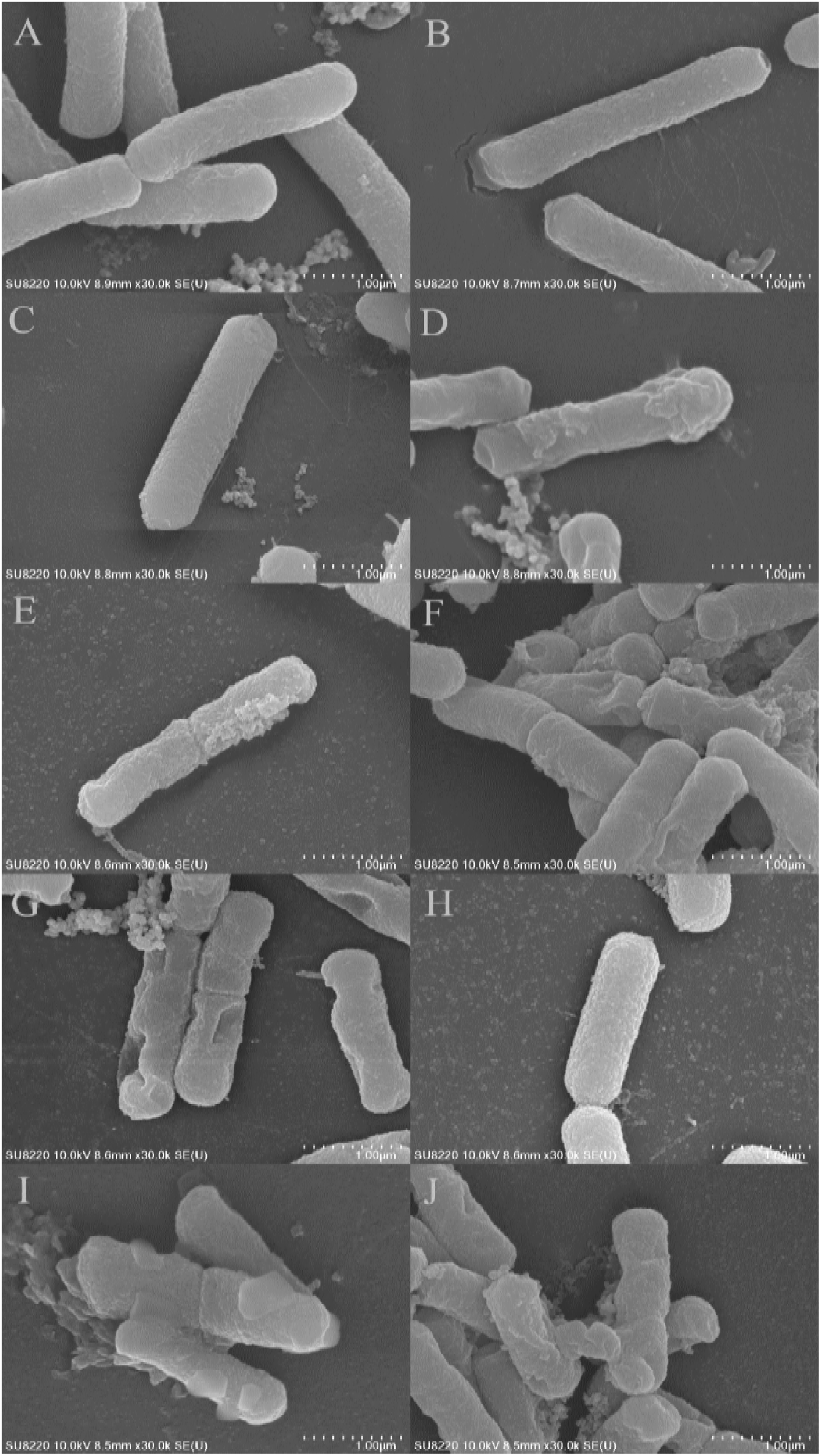
SEM images of *E. coli* after cultivation with short, medium, and long chain fatty acid monoglycerides at 37°C for 1 h. The control group (A) was not added any monoglycerides. The *E. coli* cells were treated with MB (B to D), ML (E to G), and MM (H to J) at 1/2 MIC (B, E, and H), 1 MIC (C, F, and I), and 2 MIC (D, G, and J), respectively.

#### 3.2.2 Membrane permeability measurement

The fluorescence signal of PI in *E. coli* cells was often regarded as an indicator of membrane destruction because it was captured after penetrating through damaged cell surface and binding to intracellular DNA (43). The permeability ratios of membranes of *E. coli* exposed to various concentrations of MB, ML, and MM were calculated by dividing the cell population with significant fluorescence signal by the total cell number, and the statistical results were shown in Figure 3. Compared to 25.81±1.50% of permeability ratio of *E. coli* cells in control group, the percent of cell with damaged membrane after three monoglycerides treatment all increased with an increase in concentration, but all did not exceed the MDB of 93.59±3.37%. Specifically, the monoglycerides concentrations causing above 50% of penetration ratio were 250 μg/mL for MB, 8 μg/mL for ML, and 8 μg/mL for MM, suggesting that the penetration capacity of MB to the cell membrane of *E. coli* was much lower than that of ML and MM which did not show obvious difference in penetration ability. Surprisingly, increasing further ML and MM concentration to 4 MIC led a decrease in membrane permeability, which might be due to the fact that a large part of cell death and membrane lysis appeared in membrane-damaged *E. coli* at high concentration, resulting in the appearance of the decrease trend in Figure 3 (44).

**Figure 3.**
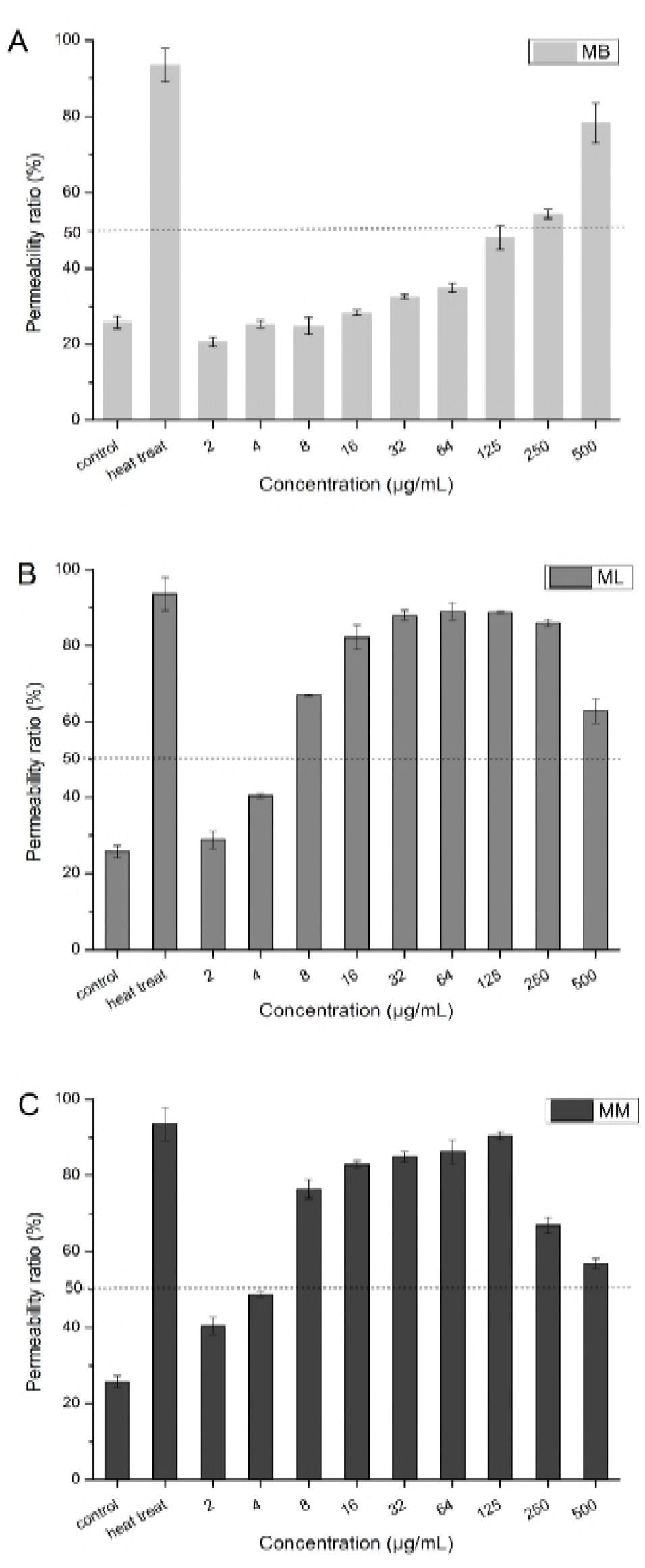
The changes in permeability ratio of membrane of *E. coli* treated with MB, ML, and MM. The statistical results of the control groups (adding the same volume of ethanol and cultivating at 37°C for 1 h), the heated treat groups (adding the same volume of ethanol and cultivating at 85°C for 1 h), and the test groups (adding MB, ML, and MM at 2, 4, 8, 16, 32, 64, 125, 250, and 500 μg/mL respectively and cultivating at 37°C for 1 h) were shown above. Error bar was stood for SD (n=3).

### 3.3 Action of monoglycerides with different chain lengths on *E. coli* DNA

#### 3.3.1 Effect of different monoglycerides on DNA structure of *E. coli*

The result of UV-visible spectral from the interaction between genomic DNA of *E. coli* and monoglycerides was shown in Figure 4. Before adding monoglyceride, the absorbance peak of *E. coli* DNA appeared at the wavelength of 260 nm. After adding three monoglycerides, the maximum absorbance values increased to varying degrees without obvious shift in wavelength, which was called hyperchromic effect, indicating that the double helix of *E. coli* DNA was destroyed by the foreign monoglycerides. Furthermore, examination the spectrum difference in MB, ML, and MM group revealed that the maximum absorbance value in MM group (0.544 at 250 nm) was greater than that in ML group (0.499 at 250 nm) and MB group (0.52 at 251 nm), suggesting that the destruction level of DNA duplexes by MM was greater than that by MB and ML. In addition, the monoglyceride concentrations causing the maximum absorption peaks in UV spectra were in great discrepancy with 16 ppm for MB, 8 ppm for ML, and 4 ppm for MM.

**Figure 4.**
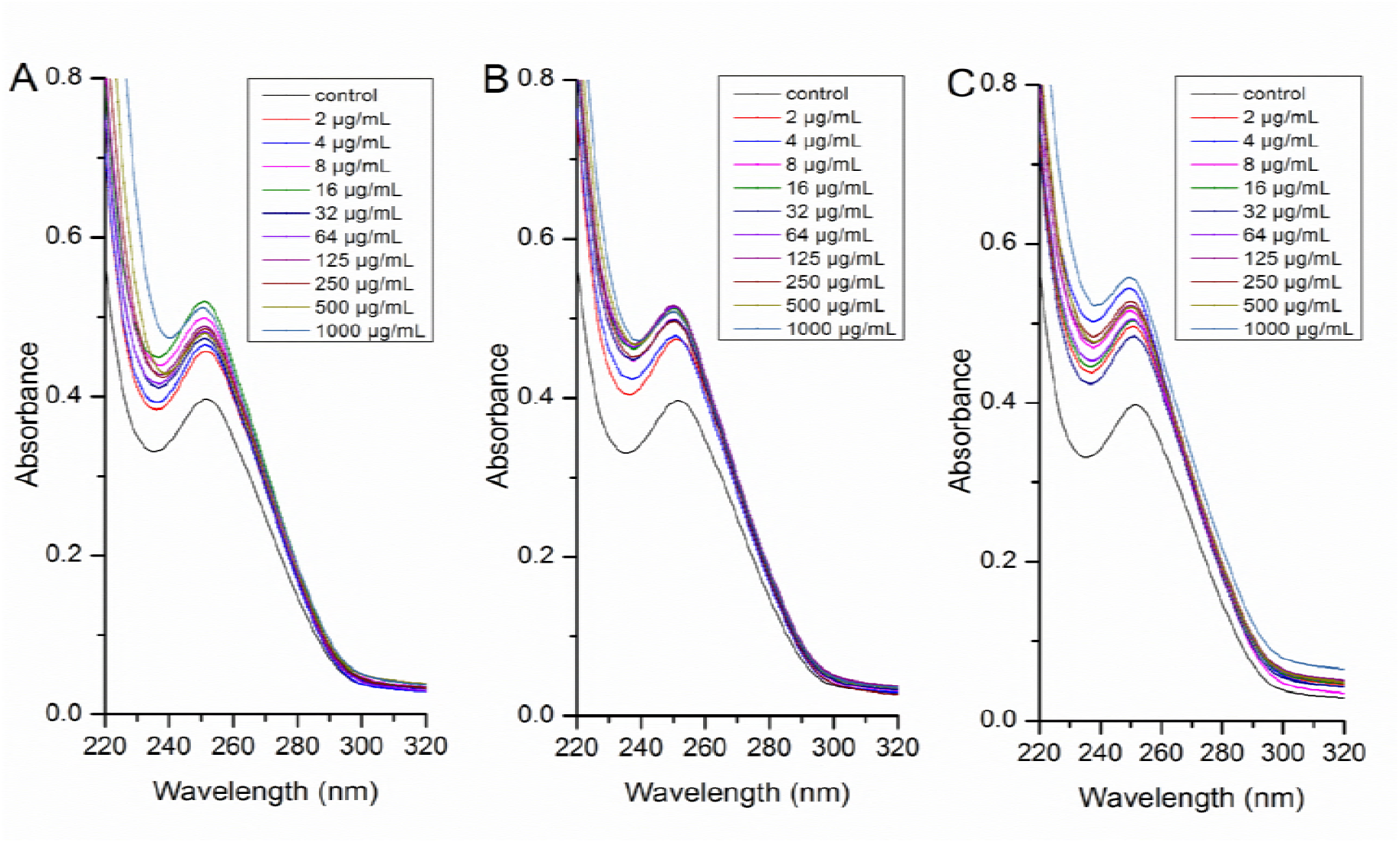
UV-visible spectrum of *E. coli* genomic DNA treated with MB (A), ML (B), and MM (C) at concentrations of 2, 4, 8, 16, 32, 64, 125, 250, 500, and 1000 μg/mL. The spectra scanning was recorded at a wavelength range from 220 nm to 320 nm, and each curve was stood for the average of three repeatly determinations

#### 3.3.2 Cell cycle changes

As shown in Figure 5, the flow histogram in control group showed a DNA distribution of a large peak close to a small peak. After adding MB, ML and MM, the two peaks in treated groups both became higher and narrower, indicating that monoglyceride treatment might alter DNA distribution in the cell cycle of *E. coli*. Furthermore, the cell proportion in G1, S, and G2 phases after the treatment of three monoglycerides was recorded in Figure 6. The G1 percents in three treated groups were all increased with different degrees, and only a significant effect was observed at 2 MIC for MB, 1 MIC for ML, and 1/2 MIC for MM. Surprisingly, further increasing MM concentration to 4 MIC resulted in a decrease in G1 proportion, which may be attributed to the leakage of DNA in damaged cells, which could be observed in Figure 2I. Comparison the discrepancy of *E. coli* cell cycle among MB, ML, and MM treatment showed the most remarkable growth appeared at MM group, followed by ML group, and MB group grew the slowest, indicating that a great discrepancy occurred at the interference action of monoglycerides with different chain lengths on the cell cycle of *E. coli*.

**Figure 5.**
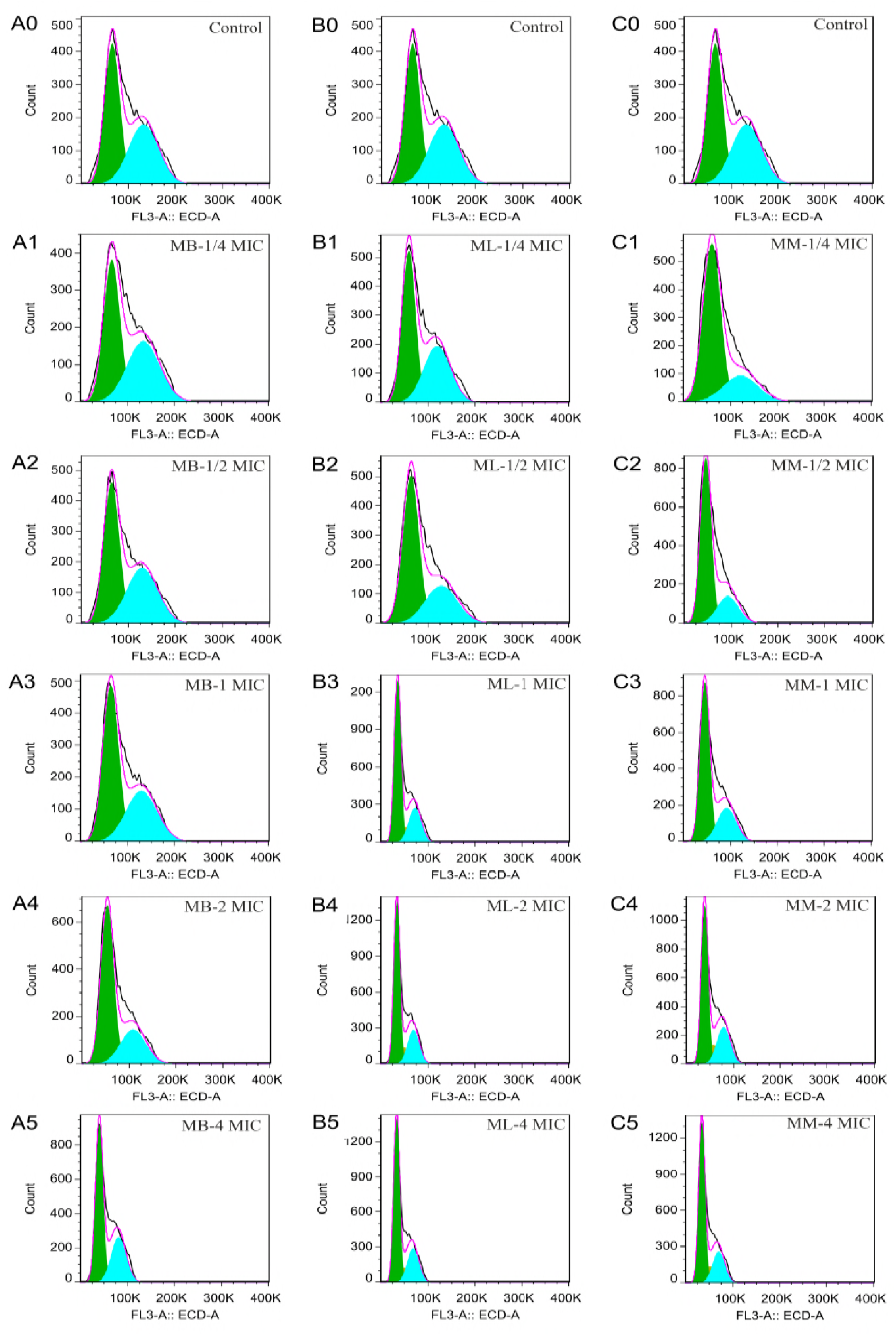
Flow histograms of *E. coli* treated with MB (A0-A5), ML (B0-B5), *and* MM (C0-C5) at 0 (A0, B0, and C0), 1/4 MIC (A1, B1, and C1), 1/2 MIC (A2, B2, and C2), 1 MIC (A3, B3, and C3), 2 MIC (A4, B4, and C4), and 4 MIC (A5, B5, and C5).

**Figure 6.**
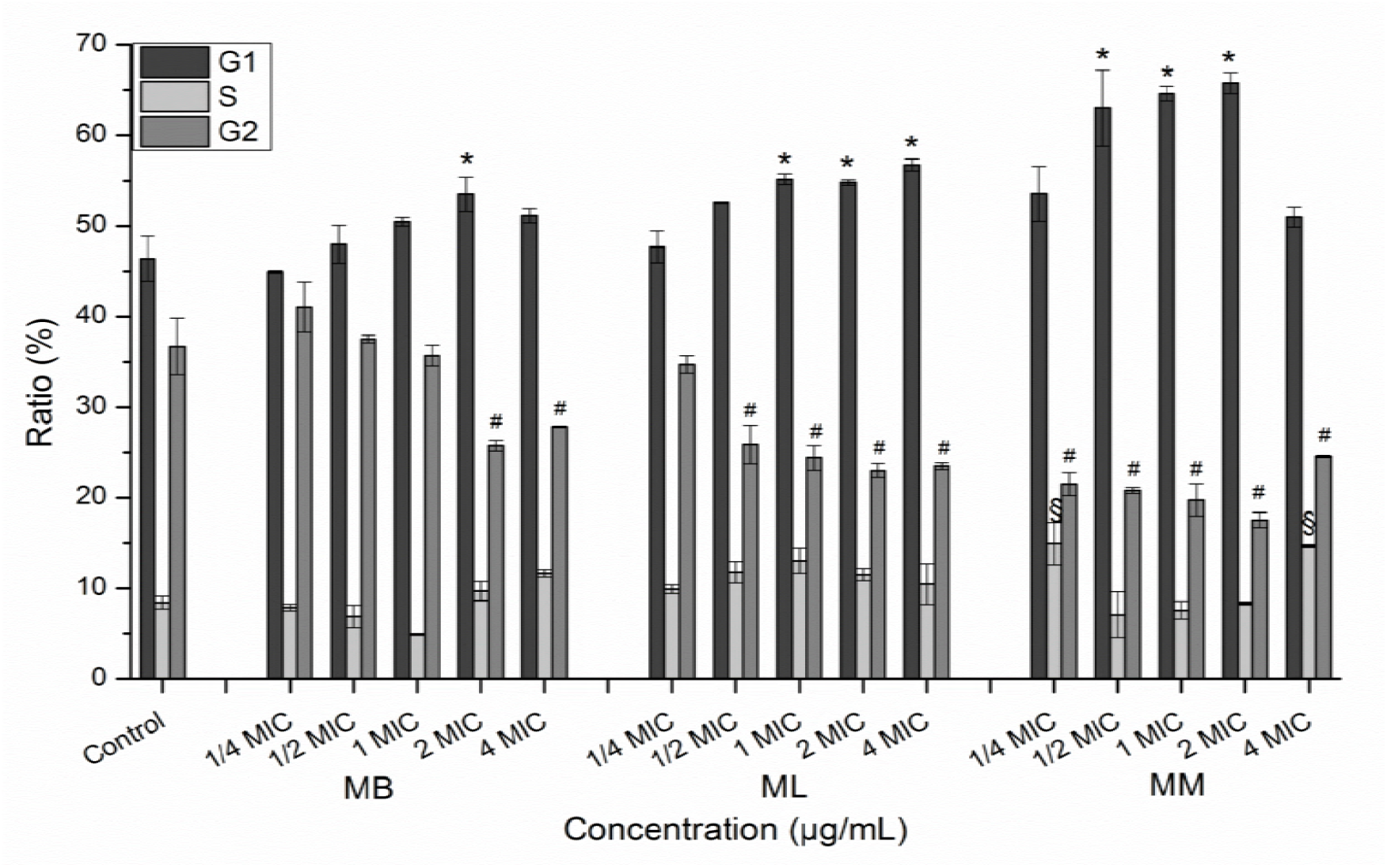
Ratio changes of G1, S, and G2 phase in *E. coli* cell cycle after treatment with MB, ML, and MM at 0, 1/4 MIC, 1/2 MIC, 1 MIC, 2 MIC, and 4 MIC. “*, $ and #” indicated statistical significant difference from the control group in G1, S, and G2 phase respectively. Error bar was represented for SD (n=3).

#### 3.3.3 Biomacromolecule synthesis inhibition

Apart from studying the effect of monoglycerides on structure and function of *E. coli* DNA, the synthesis inhibition of DNA, RNA and protein was shown in Figure 7. Compared to the steadily growth of DNA, RNA and protein content in control group, the content of the three biomacromolecules in experimental groups showed a totally different growth trend. Specifically, the RNA amount declined immediately without any time delay once adding monoglyceride, different from a surprisingly phenomenon of increasing firstly and then decreasing in DNA and protein content measurement, suggesting that RNA synthesis was firstly affected after adding monoglycerides, and then the DNA and protein synthesis was inhibited after 30min or completing a generation of cell division. Compared to timely interference in RNA synthesis, the hysteresis in the suppression of DNA and protein synthesis also implied that DNA transcription or RNA synthesis, rather than the synthesis process of protein and DNA, was the first action target in genetic central dogma. Examination the growth difference among MB, ML, and MM treatment revealed that the synthesis inhibition of ML and MM on three biomacromolecules was obviously greater than that of MB in the same conditions.

**Figure 7.**
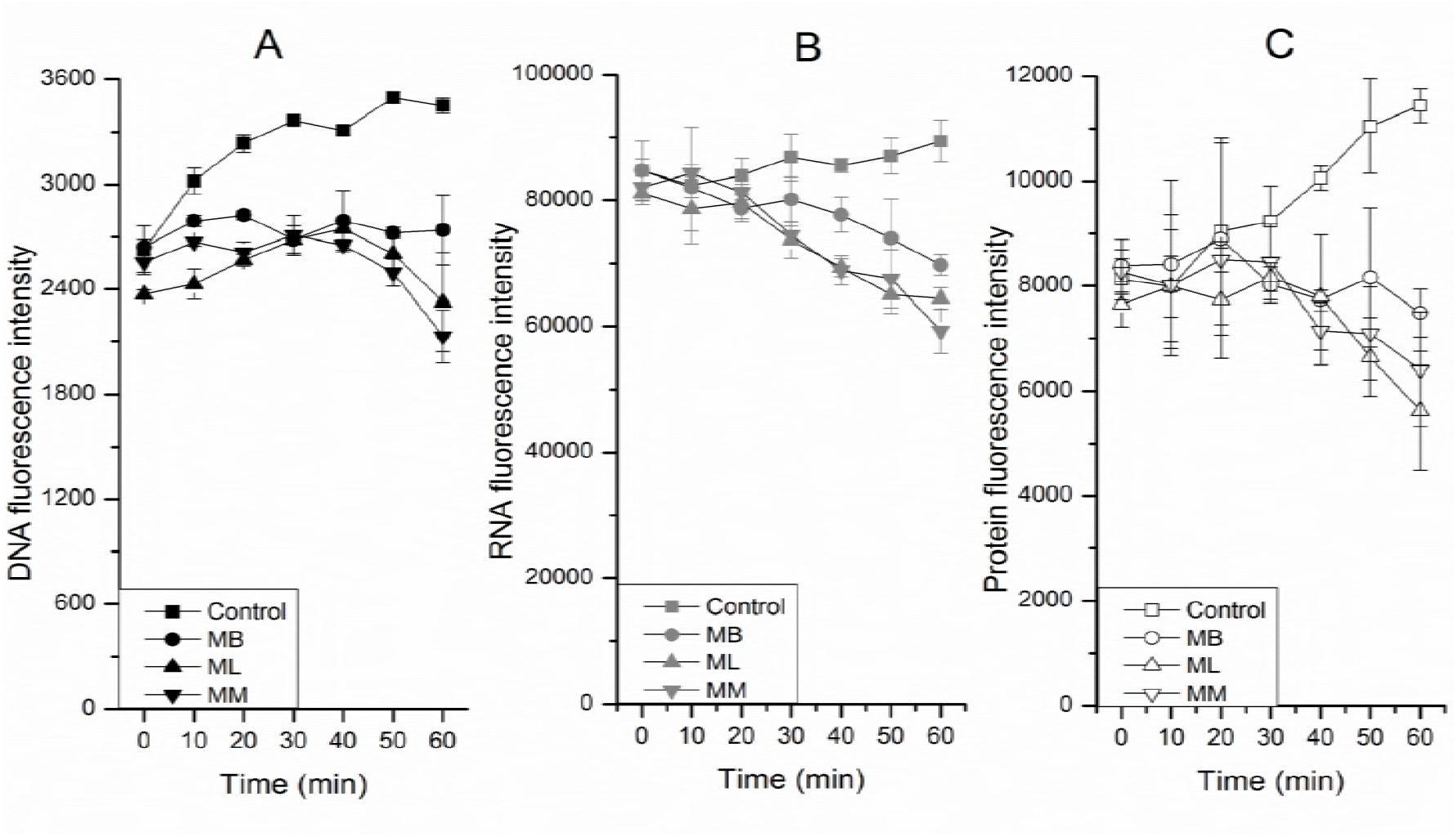
Changes of DNA (A), RNA (B) and protein (C) content in *E. coli* cell after treatment with MB, ML, and MM at MIC for 0, 10, 20, 30, 40, 50, and 60 min respectively. The control groups were added with the same volume of ethanol instead of monoglyceride solution. Error bar was represented for SD (n=3).

## 4. Discussion

### 4.1 The sensitivity of *E. coli* to the monoglycerides with different chain lengths

In general, fatty acid monoglycerides have broad antibacterial spectrum and stable inhibitory activity, which are often used to control foodborne pathogens according to some previous reports (17, 20, 21, 27, 45, 46). They offer a preliminary view on antimicrobial performance of individual monoglyceride in medium or food ingredients. Moreover, our study found that fatty acid chain length markedly affected the antibacterial performance of monoglycerides on *E. coli*, among which the sensitivity to ML and MM exceeded 8 times of that to MB. The comparative investigation in antibacterial effect of medium carbon chain monoglycerides against *E. coli* has been conducted to select the most active lipid. Clelia Altieri found *E. coli O157:H7* was completely inhibited in PC broth plus 50 μg/mL of monolaurate, whereas monomyristate and monopalmitate at the same concentration did not have the effect, which could be explained by different medium compositions and various culture conditions (26).

In this assay, the treatment with ML at 500 μg/mL or MM at 250 μg/mL killed *E. coli* cells to undetectable level, whereas MB bactericidal ability was limited even at the maximum concentration (only resulting in more than 2.2 log reduction in cell population). The huge difference in antibacterial activity among short and medium-long chain monoglycerides may be attributed to the weaker hydrophobic interaction between acyl carbon chain and the lipid membrane (47, 48). On the other hand, the hydrophobic of monoglyceride increased with increasing the length of carbon chain, which reduced its solubility in aqueous solution, thus preventing its transport through *E. coli* cell membrane (49).

### 4.2 Membrane action mechanism

At present, many studies on action mechanism of monoglyceride on bacterial cell membrane involve the change in physical morphology and the decrease in the ability to control material enter and exclude cell membrane (28, 50, 51). SEM images revealed that three monoglycerides exposure produced an uneven, rough and wrinkled surface as well as the appearance of depression on the affected *E. coli* cells, which was similar to the phenomenon reported in some literatures (21, 27, 50, 52, 53). Monoglycerides are a class of non-ionic surfactants with acyl carbon chains and hydroxyl groups, which could interact with the hydrophobic region and disrupt the composition of *E. coli* membrane (29, 54). In addition, the interaction of monoglyceride into cell membrane also increases the fluidity and permeability of the membrane (55, 56). This explanation is further supported by the growth trend of membrane permeability of monoglyceride-treated *E. coli* in Figure 3.

Review of the relationship among MIC, membrane morphology changing and cell permeability increasing revealed that the bacterial cell membrane was not the only site where monoglycerides work. As shown in Figure 2, the monoglyceride concentrations when depression and breakage appeared on *E. coli* surface were above 2 MIC (1000 μg/mL) for MB, 1 MIC (64 μg/mL) for ML, and 2 MIC (64 μg/mL) for MM, all exceeding their respective MIC, which was more like the final result of many antibacterial effects. Furthermore, the results in Figure 3 indicated that the MB, ML, and MM concentrations causing more than 50% of permeability ratio were 250 μg/mL, 8 μg/mL, and 8 μg/mL, respectively. However, compared to control groups, the three monoglycerides at the above concentrations did not cause significant decrease in cell population in experimental groups, suggesting that the increase in membrane permeability of *E. coli* had little correlation with the loss of cell viability. On the whole, the above observations showed that monoglyceride increased cell permeability at low concentrations and cause cell damage or even cell lysis at high concentrations, however, they were both not the key lethal action in the antibacterial test of monoglycerides against *E. coli.*

The reason for the difference in sensitivity to short, medium, and long chain monoglycerides may be discrepancy in action for membrane integrity. The MB concentration causing *E. coli* breakage was above 1000 μg/mL, which was far more than that of ML and MM. Similarly, the MB concentration causing above 50% of penetration ratio was 250 μg/mL, which was also much higher than that of ML and MM. The poor action effect of MB on cell membrane might be a suitable explanation for the unsatisfactory antibacterial performance in MB treatment, but was still insufficient to clarify the causes for antibacterial difference between ML and MM treatment if only in terms of the increase level in cell permeability. Considering the finding of the potential intracellular action targets in the antibacterial test of peptides (57–59), more in-depth studies are encouraged to make clear the causes for antibacterial difference of ML and MM on *E. coli*.

### 4.3 Interference of monoglyceride on the structure and function of *E. coli* DNA

There are many potential intracellular action objects after monoglyceride penetrating through cell membrane, and genomic DNA is one of the goals that cannot be ignored. According to the reports in some literatures, the modes in which foreign compounds interact with DNA are summarized as three types: intercalation binding between molecular and stacked base pairs in DNA, groove binding between molecular and major or minor grooves in DNA double-helix, and electrostatic binding between charged molecule and the external DNA backbone (60, 61). UV-visible spectral is a useful tool to study the interaction between *E. coli* DNA and foreign antibacterial agents (62). As we known, the hypochromic effect combined with a red shift in wavelength is observed only when the axial change of DNA conformation occurs, causing by the foreign molecular intercalating into DNA (63). In general, the growth magnitude in absorption spectrum is closely related to its binding force, which means the stronger its binging force, the more evident DNA hypochromic effect (64). However, Figure 4 showed a hyperchromic effect without obvious wavelength shift in UV-visible spectra, which was different from the above law of axial change, suggesting that the interaction mode between monoglyceride and *E. coli* DNA was not the typical intercalation binding. In addition, monoglyceride is an amphiphilic molecule without any charged groups, which excludes the possibility of electrostatic binding. Therefore, it is speculated that groove binding is a suitable and acceptable action mode.

The adverse impact of monoglyceride on DNA double helix may affect the DNA function and cell division, and the cell cycle is an important index for assessing whether cell division is normal or not (65). It is known that bacteria are prokaryotic cells that have phases I, R, and D in the cell cycle, corresponding to phases G1, S, and M in eukaryotic cells (66). Unlike eukaryotic cells, bacteria will directly enter into the phase D to begin chromosome separation immediately without going through phase G2. Flow cytometry is a common method to investigate the effect of foreign substances on the cell cycle. Figure 5 showed that peak shape in flow histogram of *E. coli* was obviously affected by MB, ML, and MM. Specifically, the growth in phase G1, and the decline in phases S and G2 in Figure 6 indicated that monoglyceride firstly disturbed phase G1 instead of phases S and G2, causing *E. coli* cells to arrest in phase G1 and therefore were unable to complete normal cell division. The finding does not fully correspond to some previous reports of antibacterial agents (67, 68). PAN Ling Zi also found that phase G1 of cell cycle was evidently affected in *GS115* after melittin treatment for 4h (66). Whereas, Jin Cai revealed that the Laminaria japonica extract acted on phase R (or phase S) rather than phase I (or phase G1), causing cell cycle of *C. michiganense* arrest in phase R (69).

In order to further study the correlation between DNA double helix disruption and abnormal cell cycle, the synthesis of intracellular DNA, RNA, and protein was evaluated using flow cytometry. The results presented in Figure 7 indicated that DNA, RNA, and protein synthesis are affected after adding monoglyceride, and the time of DNA and protein synthesis receiving inhibition are much later than that of RNA. Similar studies have been conducted to figure out the effect of antimicrobial peptides on bacterial macromolecular synthesis (70–72). Aleksander Patrzykat found that the RNA synthesis in *E. coli CGSC 4908* was affected after adding pleurocidin peptide at MIC within 5 min, followed by the suppression for DNA synthesis, whereas protein synthesis was not obviously affected except for a decrease in growth rate (73). Evan F. Haney also observed a similar growth trend of DNA, RNA, and protein synthesis in *E. coli CGSC 4908* with three Puro B peptides treatment for 30 min (67). The difference between the assay and previous reports is due to differences in antibacterial agent types, binding modes with DNA and action time (72, 74, 75). Combination with the action site of monoglyceride on the cell cycle of *E. coli*, it is speculated that DNA double helix is firstly damaged, followed by the inhibition of RNA synthesis, subsequently protein and DNA synthesis is also affected, resulting in DNA replication suppression and cell cycle arrest in phase G1.

A new concentration-dependent antibacterial mechanism is summarized in Figure 8. The monoglyceride firstly crosses *E. coli* membrane at concentrations far below the MIC, which is the basis for the antibacterial effect. Then it targets to genomic DNA and damages its double helix, affecting RNA synthesis and subsequent protein and DNA synthesis, causing cell cycle arrest, and finally resulting in cell division disorder. The interference of cell cycle often means the inhibition of cell division which occurred at concentrations close to MIC; therefore, we consider this type of intracellular action as the key lethal effect in the antibacterial test of monoglycerides on *E. coli.* And the membrane lysis and cell disruption appear when adding monoglycerides at 2 MIC or more, which is more like the final result of a combination of various antimicrobial modes.

**Figure 8.**
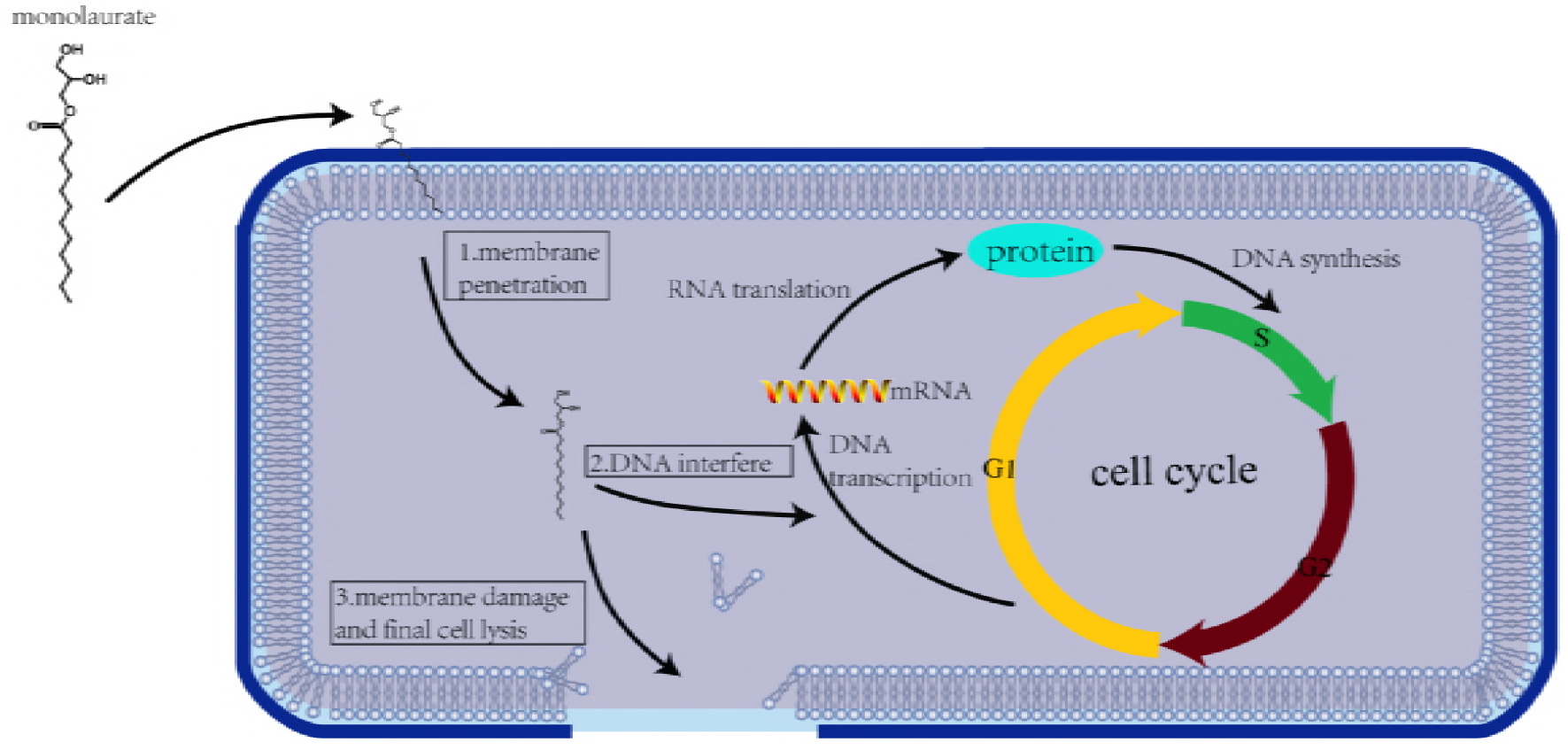
Explanation of the overall inhibitory mechanisms of glycerol monoglyceride on *E. coli* with monolaurate as an example. ML firstly crossed the cell membrane and interfered with the normal function of the DNA, eventually leading to cell lysis. The action site of ML on DNA was identified as the process of DNA transcription, causing the reduction in the synthesis of RNA and protein, resulting in cell cycle arrest and ultimately cell division inhibition.

The differentiated performance of ML and MM treatment in UV-visible spectra and cell cycle assay may be used for explaining the causes for different sensitivity of *E. coli* to two monoglycerides. The saturation action concentration of ML in hyperchromic effect was 8 μg/mL, which was higher than 4 μg/mL of that in MM treatment. Furthermore, a significant growth in phase G1 after adding ML was observed at 64 μg/mL, much larger than 4 μg/mL of concentration observed in MM treatment. These data revealed that the saturated action concentration of MM to DNA double helix and cell cycle in *E. coli* was obviously smaller than that of ML, indicating that the inhibitory effect of MM to *E. coli* DNA was greater than that of ML at doses near MIC. As for the reason for the different affinity between two esters and *E. coli* DNA, we are still not clear, which may need to be further studied by the interaction force including hydrogen bond, van der Waal’s force, and hydrophobic interaction (76, 77).

In conclusion, the study presented here explained the cause for the divergence in the antibacterial activity of short, medium and long chain monoglycerides on *E. coli*, and identified DNA interference and cell cycle arrest as the key lethal effect in the antibacterial behavior of monoglyceride, and proposed a new dose-dependent inhibitory mechanism. In three monoglycerides, MB has the worst antibacterial effect due to its poorest membrane permeability, and the superiority of MM in comparison with ML is mainly due to its stronger DNA destroying ability and higher interference effect on cell cycle, which also means better inhibition ability on cell division. As for the inhibitory modes, the increase of membrane permeability at low doses is the basis for playing the antibacterial effect, then the arrest of cell cycle at doses near MIC is the key lethal effect in antibacterial test, and the final cell lysis at high doses is considered as the result of synergistic action of various antibacterial effects. In addition, a possible route was proposed to explain the correlation between DNA double helix disruption and cell cycle arrest.

## Acknowledgments

Mr. Hao Cheng and Mr. Tao Ruan are acknowledged for their assistance in operating flow cytometer and scanning electron microscope. Dr Yaping Nie is greatly appreciated for her useful suggestion in the aspect of cell cycle.

## Table of contents graphics

The main content of the manuscript is summarized in a figure. Glycerol monoglyceride with different chain lengths shows different biological activity against *Escherichia coli* and their antibacterial mechanism can be summarized as the following: the antibacterial basis is the increase of membrane permeability at low concentration, the key action is the inhibition of cell division in medium concentration near MIC, and the cell lysis at high concentration can be regarded as the result of combination action of multiple antibacterial effects.

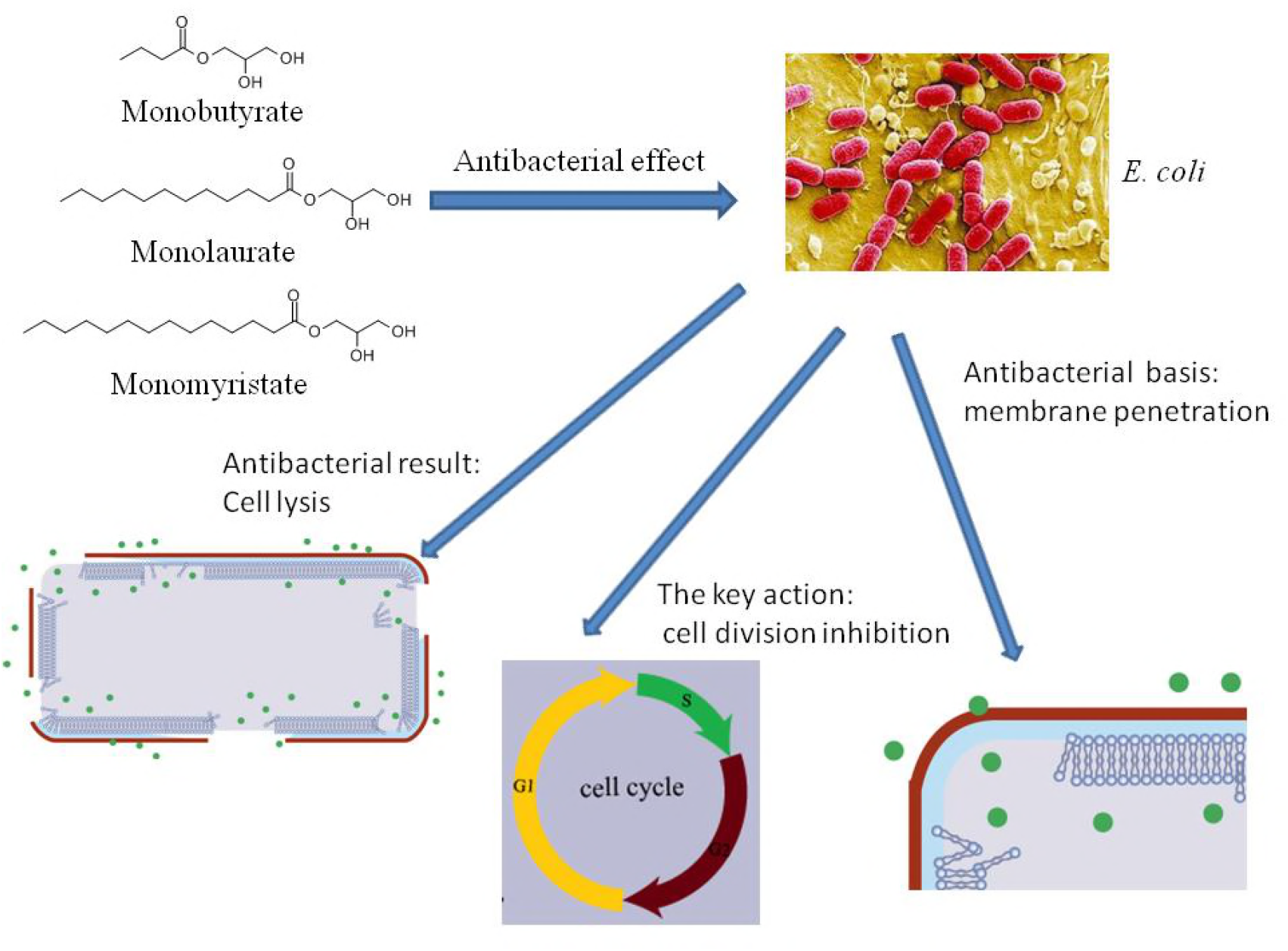

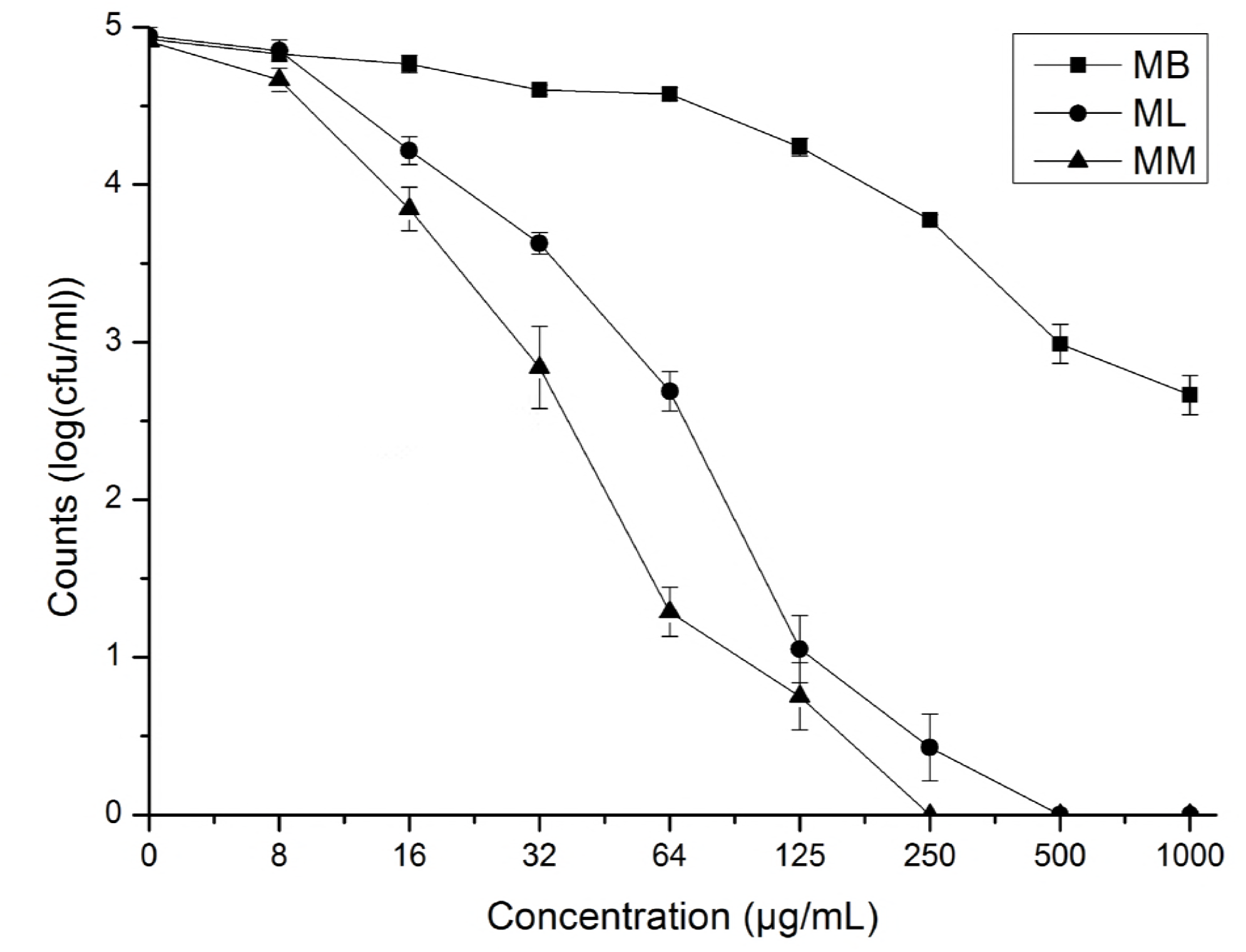

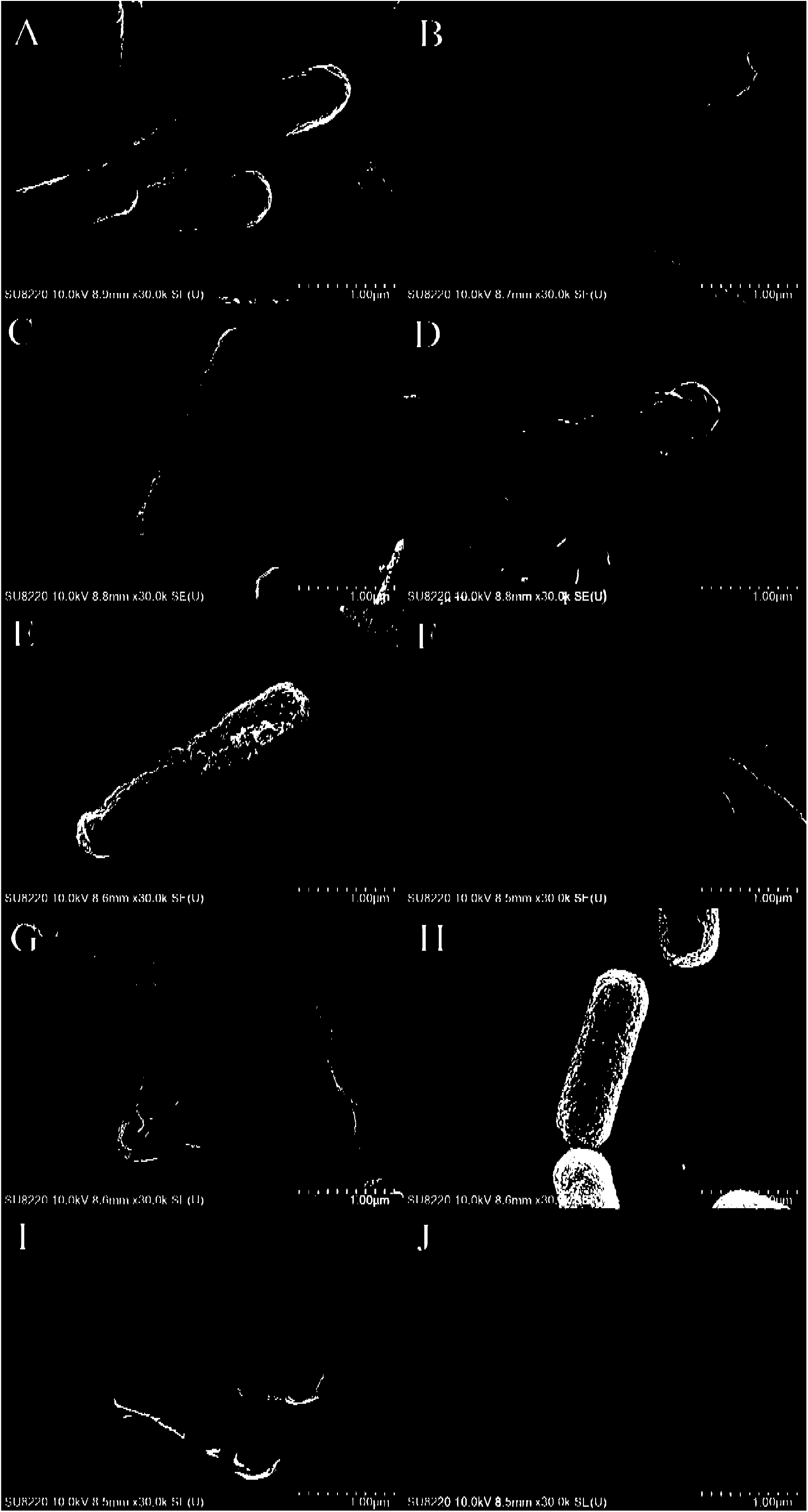

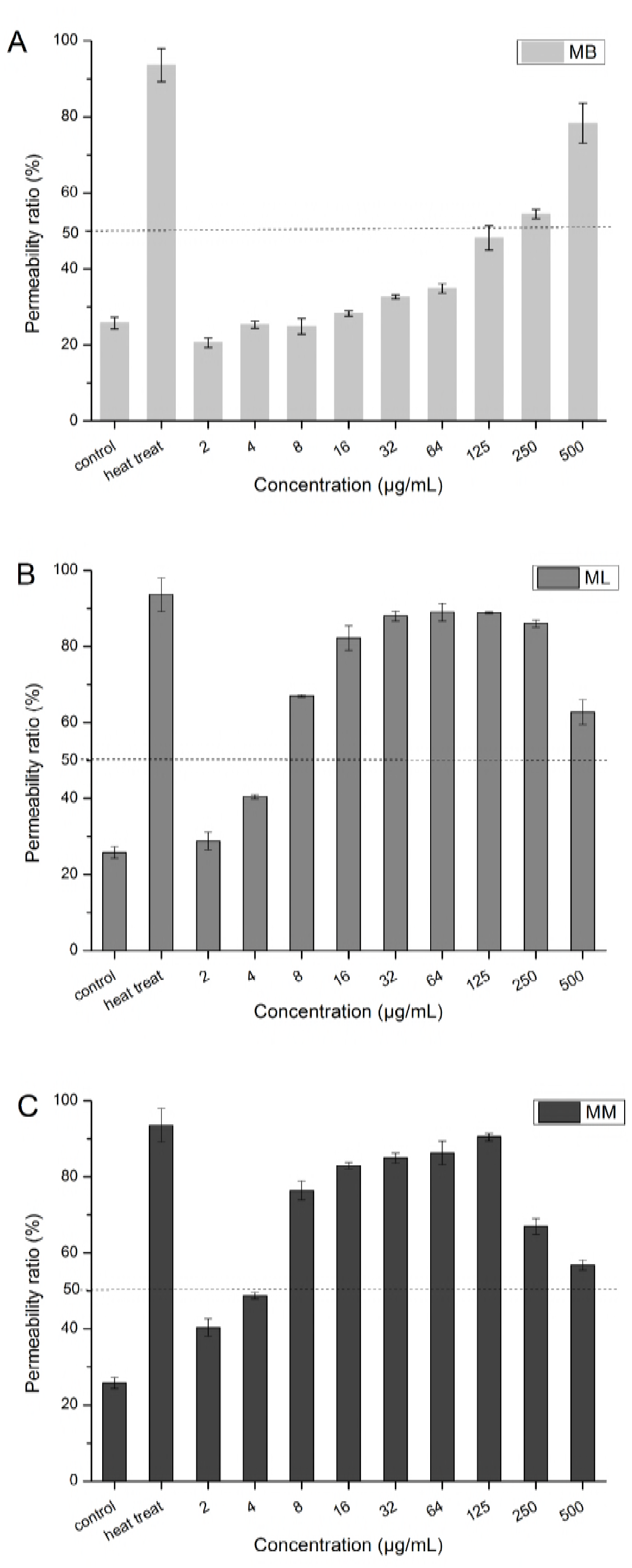

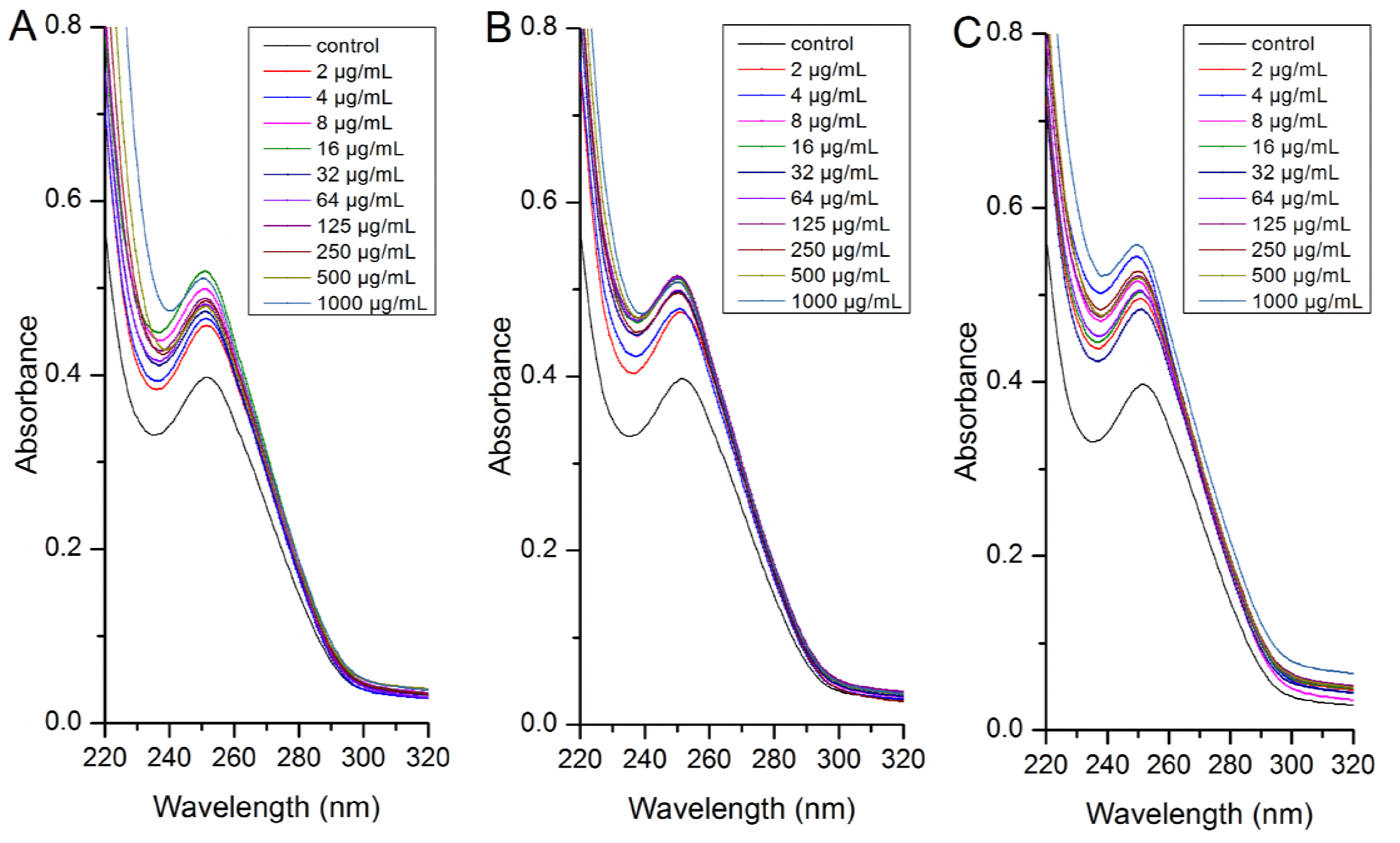

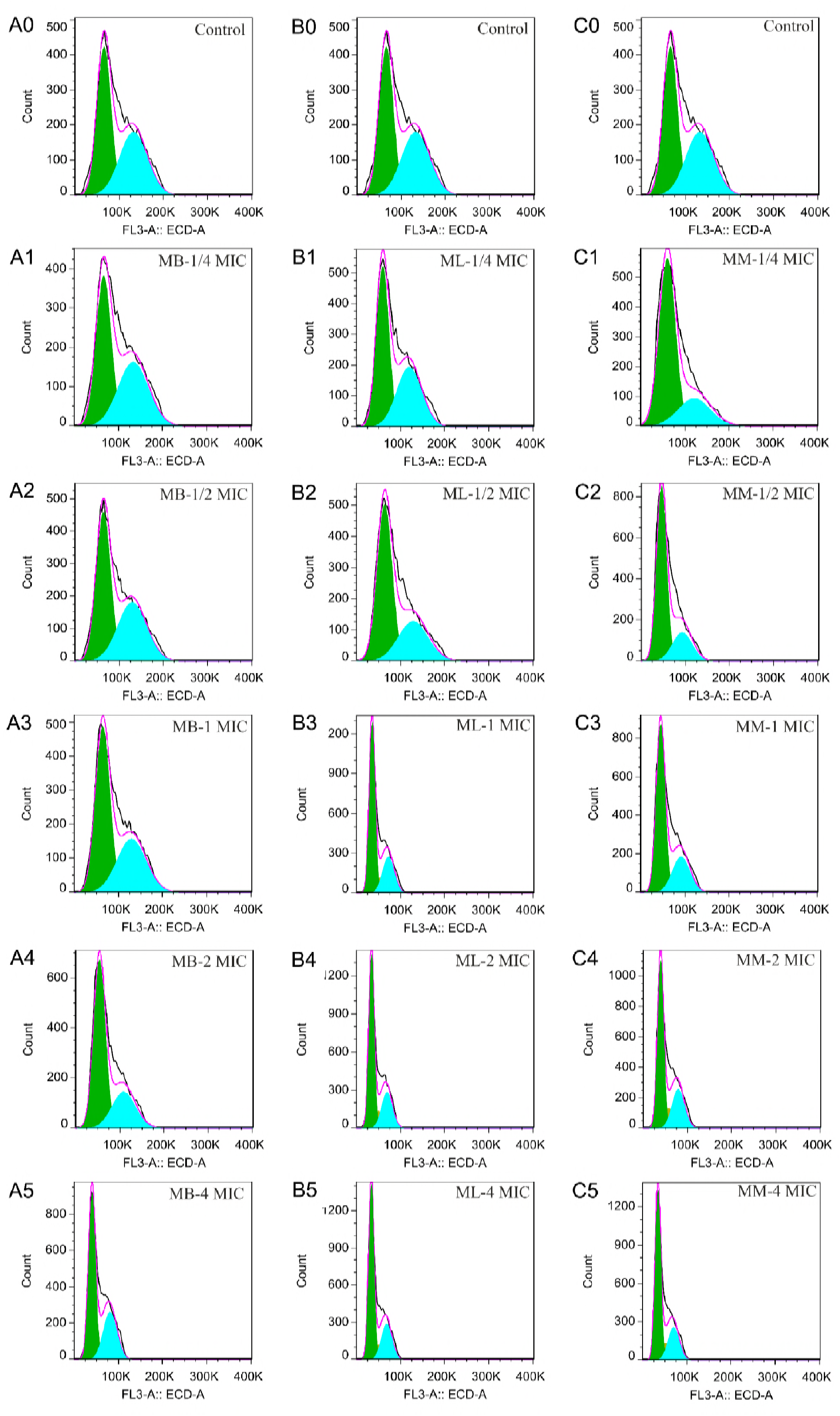

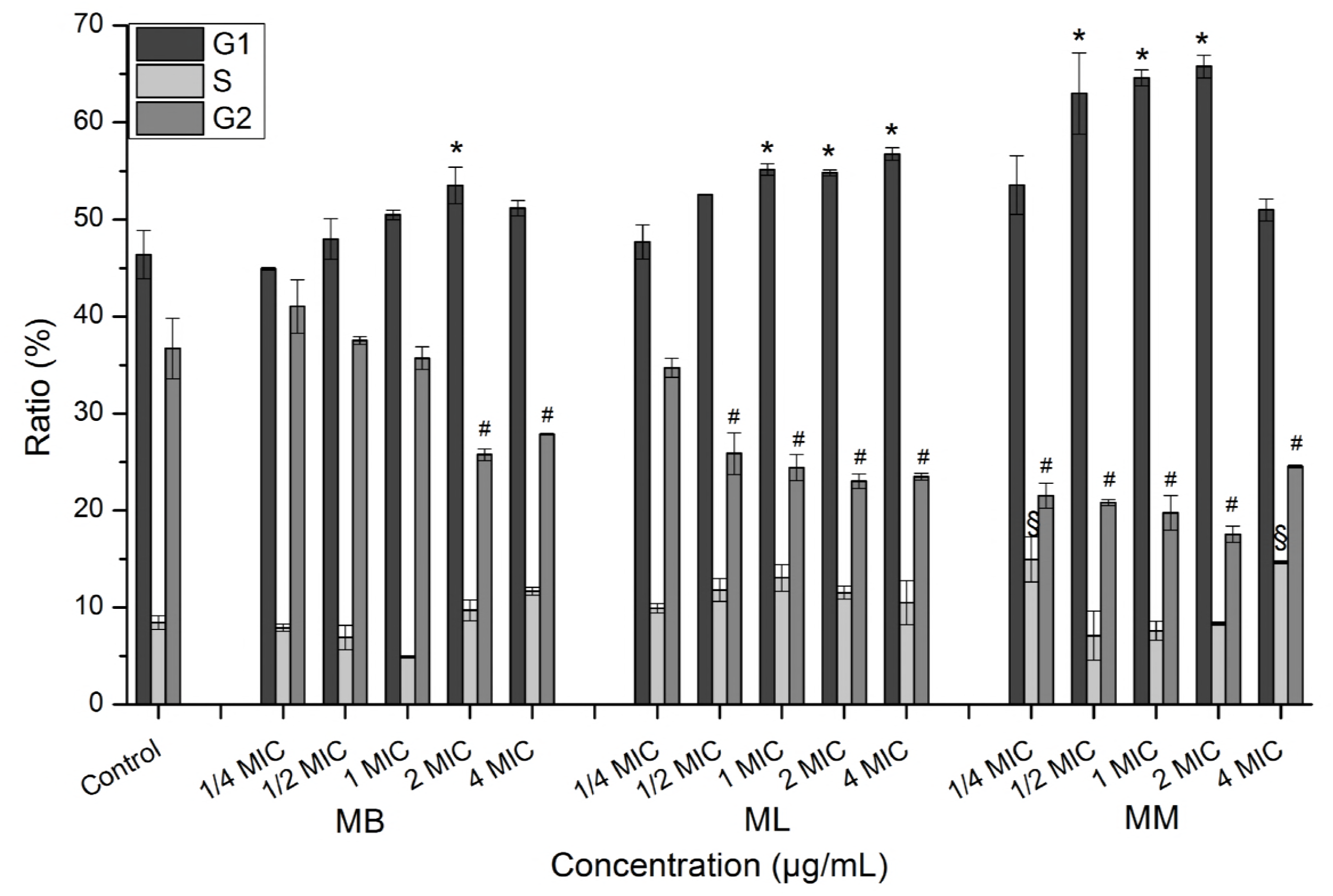

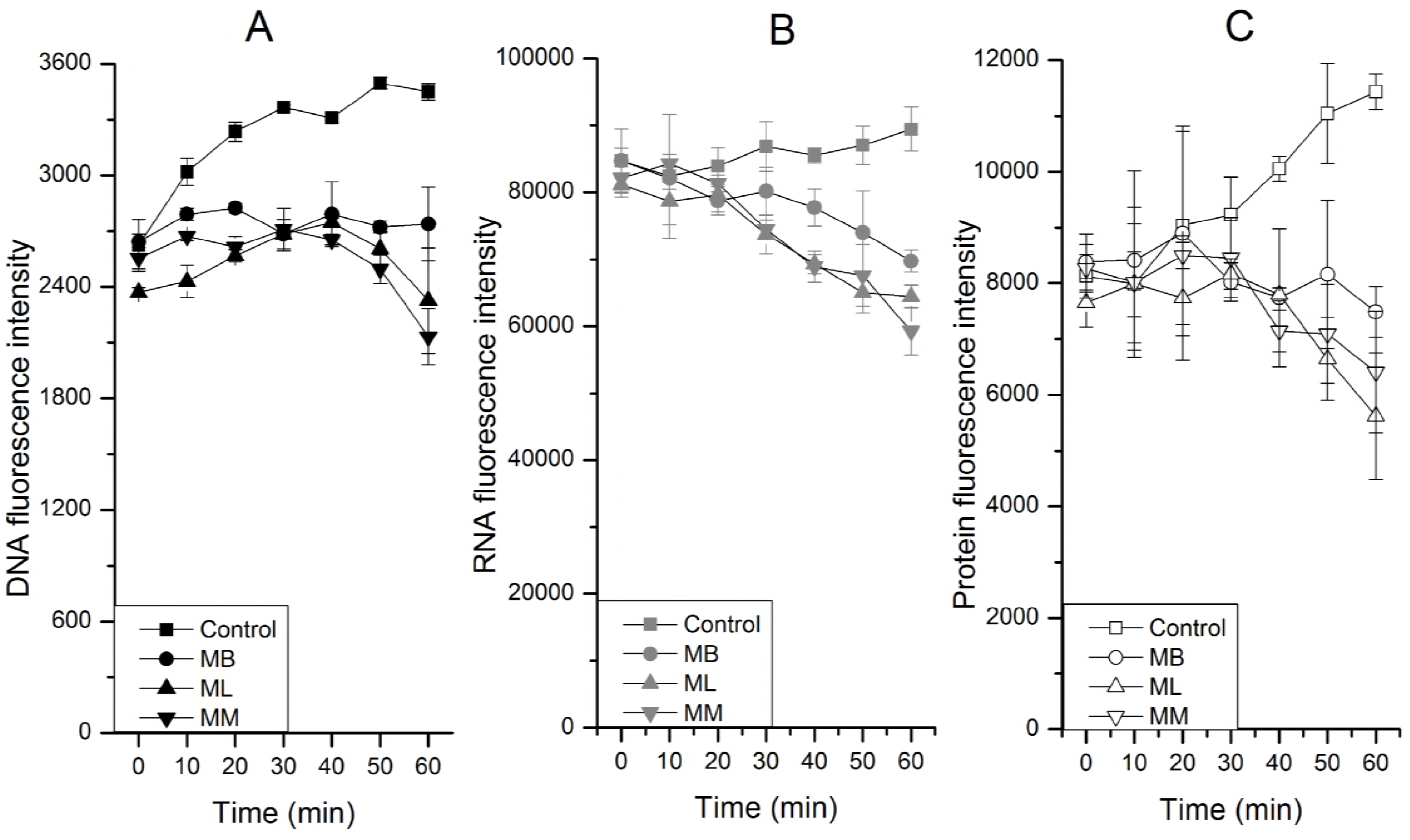

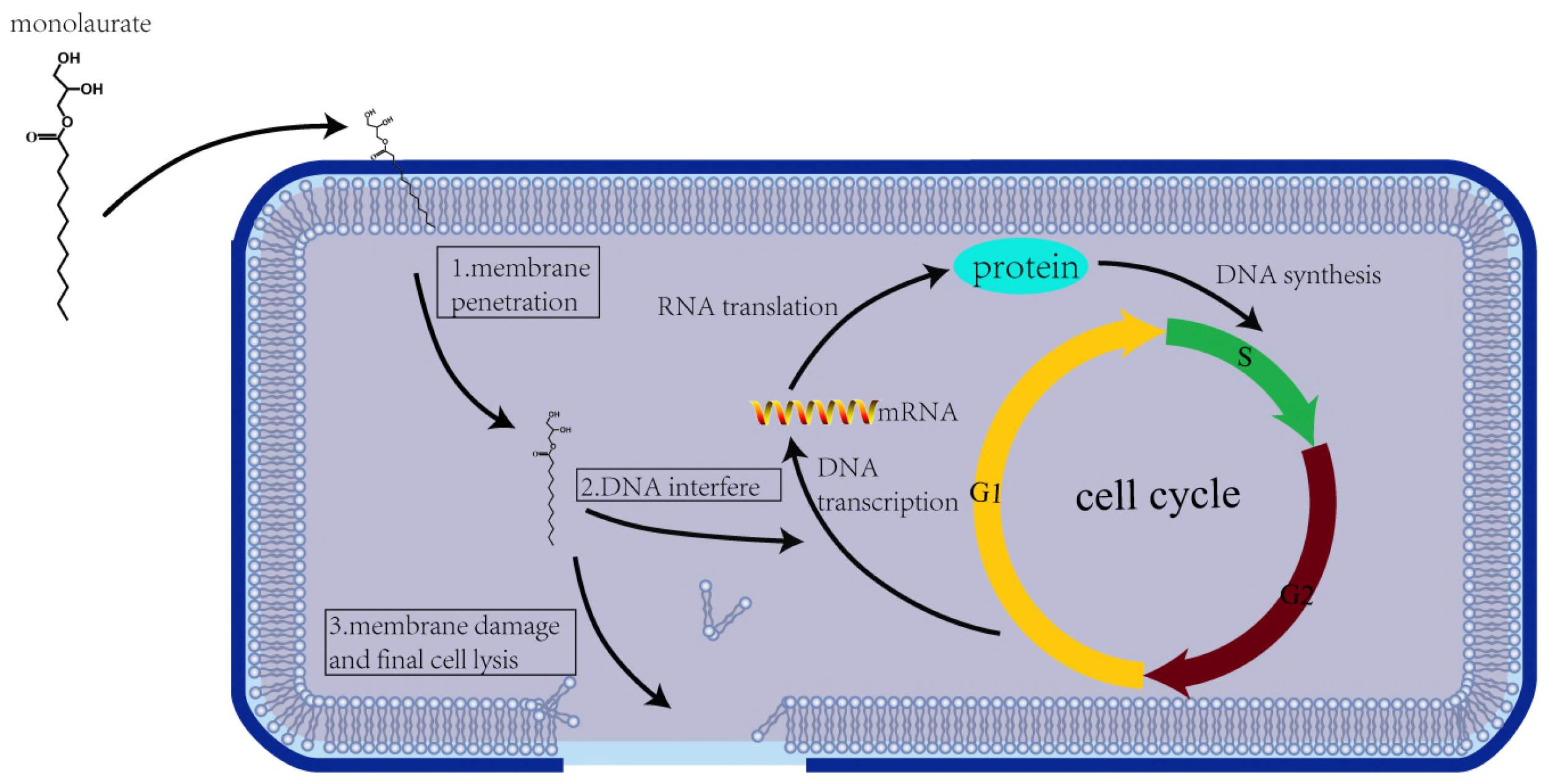

